# Primary Ciliary Dyskinesia patient specific hiPSC-derived airway epithelium in Air Liquid Interface culture recapitulates disease specific phenotypes *in vitro*

**DOI:** 10.1101/2023.04.25.538316

**Authors:** Laura von Schledorn, David Puertollano Martín, Nicole Cleve, Janina Zöllner, Doris Roth, Ben Ole Staar, Jan Hegermann, Felix C. Ringshausen, Janna Nawroth, Ulrich Martin, Ruth Olmer

## Abstract

Primary ciliary dyskinesia (PCD) is a rare heterogenic genetic disorder associated with perturbed biogenesis or function of motile cilia. Motile cilia dysfunction results in diminished mucociliary clearance (MCC) of pathogens in the respiratory tract and chronic airway inflammation and infections successively causing progressive lung damage. Current approaches to treat PCD are symptomatic, only, indicating an urgent need for curative therapeutic options. Here, we developed an *in vitro* model for PCD based on human induced pluripotent stem cell (hiPSC)-derived airway epithelium in Air-Liquid-Interface cultures. Applying transmission electron microscopy, immunofluorescence staining, ciliary beat frequency and mucociliary transport measurements, we could demonstrate that ciliated respiratory epithelia cells derived from two PCD patient specific hiPSC lines carrying mutations in *DNAH5* and *NME5*, respectively, recapitulate the respective diseased phenotype on a molecular, structural and functional level.

## 1. Introduction

Motile cilia, membrane-bound organelles, are involved in developmental processes and function of different organs. They are present on various cells, e.g., in the ventricular system of the brain, on nodal cells during development that are responsible for left-right asymmetry, in the fallopian tubes, and the epithelium of the respiratory tract [1]. Mutations in genes involved in assembly and function of motile cilia can lead therefore to complex disorders summarized as motile ciliopathies [1]. Primary ciliary dyskinesia (PCD) is a rare genetic disorder belonging to the group of motile ciliopathies, with current rates estimated at 1:10.000 inhabitants depending on the degree of consanguinity within a particular population [2, 3]. PCD is a mainly autosomal recessive and very rarely X-chromosomal recessive or autosomal dominant condition and up to date PCD-causative mutations in more than 50 genes are known, leading to different functional defects ranging from abnormal beat pattern through impairment of dynein arms to a complete absence of motile cilia [3, 4]. Patients with PCD suffer from dysfunction of the cilia in the respiratory epithelium of the upper and lower respiratory tract, leading to impaired mucociliary clearance (MCC), chronic inflammation and infection of the airways and the lung. In addition, depending on the underlying mutation, laterality defects, congenital heart disease and infertility or subfertility are further symptoms of PCD [4]. So far, PCD research is mainly focusing on identification of disease causing mutations and affected genes by utilizing primary airway or nasal epithelial cells from patients with PCD [5]. While in particular nasal epithelial cells are generally accessible from PCD patients, these cells have important disadvantages, e.g., the small number of cells within obtained samples, the difficulties to obtain samples in regular intervals, bad sample quality due to pre-damaged tissue and the missing expandability under cell culture conditions. Clearly, other cell sources are required to develop advanced organotypic culture models. Human induced pluripotent stem cells (hiPSCs) can be generated from patients’ somatic cells [6-10], for instance from hair or small blood samples, are generally considered as an excellent tool for modelling genetic diseases in vitro and have already been used for drug screening [11, 12]. Established protocols for mass expansion of undifferentiated hiPSCs as well as upscaling of differentiation cultures allow for generation of clinically relevant cell numbers [13-17]. In contrast to primary airway cells, hiPSCs allow for clonal genetic engineering by innovative gene editing techniques like TALENs or CRISPR/Cas9 and therefore seamless correction of e.g. disease underlying point mutations [18] or introduction of disease relevant mutations [19]. Genetic diseases have already been modelled using patient specific hiPSCs, e.g., pulmonary aterial hypertension[20], or cystic fibrosis (CF). In the case of CF, we and others have previously shown the generation of patient-specific hiPSCs [18, 21], their genetic correction *in vitro* and application of CF-hiPSC-derived epithelial cells for High-Throughput (HT) screening to identify new modulators or drug testing of Cystic Fibrosis Transmembrane Conductance Regulator (CFTR) function [11, 12, 22]. In the case of PCD, two reports applied hiPSCs to further explore the disease [23, 24]. While analyses of Hawkins et al. [23] just showed that similar to mutated primary airway epithelia *DNAH5*-mutated hiPSC-derived epithelial cells do not express the DNAH5 protein and consequently carry non-motile cilia with a reduced number of outer dynein arms, Sone et al. recently described the modelling of ciliary function using hiPSC-derived epithelial cells in more detail [24]. Since it has been difficult to recapitulate unidirectional mucociliary flow using hiPSC-derived airway epithelium *in vitro*, Sone et al. utilized a complex airway on a chip technology to demonstrate that abnormal mucociliary transport and altered ciliary ultrastructure, both hallmarks of PCD that are detectable in primary material, are also detectable in PCD patient-derived hiPSC with different underlying mutations (HEATR2, DNAH11 and PIH1D3) [24]. Here we show, that also a conventional Air-Liquid-Interface (ALI) culture system, which is readily available in many laboratories, facilitates investigation of cilia structure and function in hiPSC-derived ciliated airway epithelial cells. Absence of ciliary movement or altered ciliary beating, respectively, and reduced particle transport as surrogate for MCC, as well as altered cilia ultrastructure compared to hiPSC-derived WT airway cells could be confirmed in PCD patient hiPSC-derivatives carrying mutations in *DNAH5* [9], encoding for a protein of the outer dynein arm, and NME5 [7], encoding for a protein of the radial spoke.

## 2. Results

### PCD specific hiPSC lines can be efficiently differentiated towards respiratory epithelial cells

For establishment of a PCD in vitro disease model, two hiPSC lines from patients carrying homozygous mutations in *DNAH5* (c.7915C > T [p.Arg2639^*^]; hereafter DNAH5^mut^ Clone 22 and Clone 24) and *NME5* (c.415delA [p.Ile139Tyrfs^*^8]; hereafter NME5^mut^), respectively, were previously generated [10]. A non-PCD diseased hiPSC line (non-PCD) was used as control [25].

In order to direct the hiPSCs towards respiratory airway epithelial cell fate, a stepwise differentiation protocol was used that induced definitive endoderm (DE), anterior foregut endoderm (AFE) and lung progenitor cells (Fig. 1 A). The protocol was slightly adapted individually for all cell lines regarding seeding densities and duration of AFE induction.

**Figure 1:**
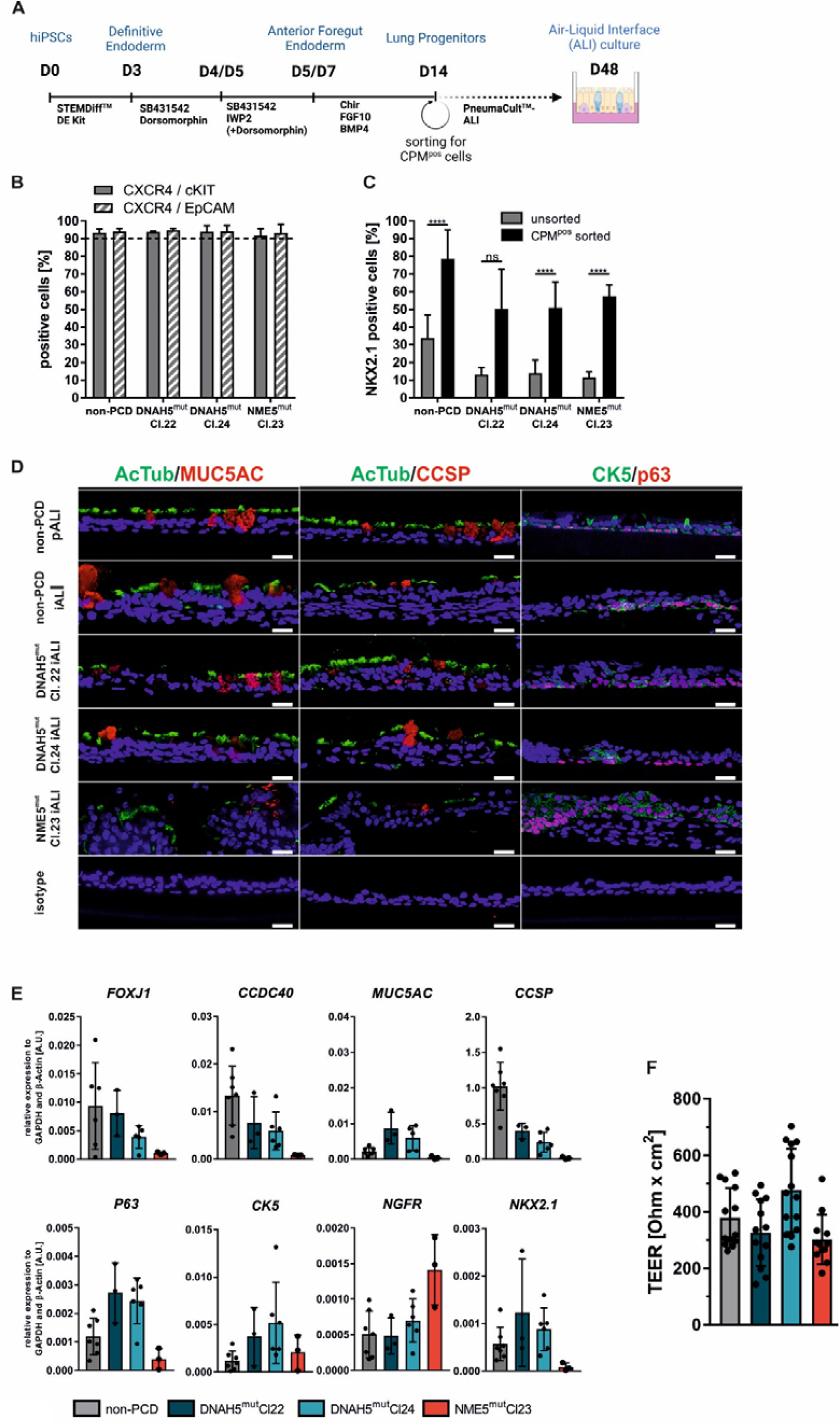
PCD specific hiPSC lines can be efficiently differentiated towards ciliated respiratory epithelium. Schematic depiction of the stepwise differentiation protocol for generating respiratory airway epithelium from human induced pluripotent stem cells (hiPSCs). Scheme generated with Biorender.com (A). Differentiation efficiencies of definitive endoderm (DE) induction measured by flow cytometry staining of CXCR4, cKIT and EpCAM (means ± SD, n=3-23 independent experiments) (B). Proportion of NKX2.1 positive cells at day 14 of differentiation before (unsorted) and immediately after MACS (CPM^pos^ sorted) (means ± SD, n=3-19 independent experiments) (C). Immunofluorescence staining of paraffin embedded ALI culture cross-sections of indicated cell lines at day 28 after air lift. Immunolabeling with acetylated tubulin (AcTub) and MUC5AC (left panel), AcTub and CCSP (middle panel) and p63 and CK5 (right panel); all scale bars represent 20 μm (D). mRNA expression of airway epithelial markers (ciliated cells: *FOXJ1, CCDC40;* goblet cells: *MUC5AC;* club cells: *CCSP;* basal cells: *P63, CK5, NGFR*; lung progenitor cells: NKX2.1) in hiPSC-derived ALI cultures at day 26-32 after air lift (means ± SD, n=3-7 independent experiments) (E). Transepithelial electrical resistance (TEER) measurements of different hiPSC-derived ALI cultures at day 26-29 after air lift. Each point represents an individual insert (means ± SD, n=3-4 independent differentiations) (F).

By applying these cell-line-specific optimized conditions, more than 91% of the cells expressed the definitive endoderm specific markers CXCR4, cKIT and EpCAM at day 3 of differentiation in all hiPSC lines (Fig. 1 B). Further differentiation to the respiratory lineage resulted in expression of the lung progenitor cell specific transcription factor NKX2.1 in 11-33 % (non-PCD: 33.7±13.1%; DNAH5^mut^ Cl. 22: 13.1±4.1%; DNAH5^mut^ Cl. 24: 14.0±7.3% and NME5^mut^: 11.6±3.1%) of the cells at day 14 of differentiation. In order to enrich lung progenitor cultures, magnetic activated cell sorting (MACS) using the cell surface maker carboxypeptidase M (CPM) was performed [26]. Sorting allowed significant enrichment of lung progenitor cells to 50-78% NKX2.1 expressing cells (non-PCD: 78.5±16.5%; DNAH5^mut^ Cl. 22: 50.2±22.4%; DNAH5^mut^ Cl. 24: 50.8±14.7%; NME5^mut^: 57.4±6.4%) (Fig. 1 C).

For maturation towards respiratory airway epithelium, sorted lung progenitor cells were seeded onto transwells and cultivated under ALI conditions for at least 28 days before analysis. Immunofluorescence staining of ALI cross-sections shows presence of airway epithelium specific cell types (ciliated cells (AcTub), goblet cells (MUC5AC), club cells (CCSP) and basal cells (P63 and CK5)) in all cell lines (Fig. 1 D). Remarkably, NME5^mut^-derived cultures generated thicker cell layers and less AcTub, MUC5AC and CCSP positive cells as compared to DNAH5^mut^ and non-PCD diseased counterparts. Primary bronchial epithelial cell (PBEC)-derived ALI cultures served as control (non-PCD pALI). Additionally, airway epithelial cell specific marker expression was confirmed by RT-qPCR (Fig. 1 E). Expression levels of all tested genes in DNAH5^mut^ ALI cultures were comparable to those of non-PCD disease cultures. Although all markers were detectable, NME5^mut^ cultures showed lower expression for most tested markers. To assess epithelial integrity, transepithelial electrical resistance (TEER) measurements were performed after 28 days of ALI culture and showed TEER values of 302-476 Ωxcm^2^, indicating intact and comparable barrier function in cultures derived from all cell lines (Fig. 1 F).

### PCD specific hiPSC-derived epithelial cells show impaired expression of cilia proteins

In order to visualize ciliary protein expression, ALI cultures at day 28 post air lift were dissociated into single cells and stained for DNAH5 and NME5 protein, respectively. Immunofluorescence staining against AcTub was used to identify cilia and confirmed the presence of cilia with normal appearance in all cell lines. In ciliated cells generated from non-PCD diseased primary cell-derived ALI cultures (non-PCD pALIs; Fig. 2 A, B) as well as non-PCD hiPSC-derived ALIs (non-PCD iALIs; Fig. 2 C, D), expression of DNAH5 and NME5 was observed that was associated with the cilia. In contrast, DNAH5-mutant ciliated cells (DNAH5^mut^ Cl. 22 and Cl. 24) showed absence of DNAH5 protein in the cilia. NME5 protein expression remains unchanged in these cells (Fig. 2 E-H). In ciliated cells derived from NME5-mutant hiPSCs, cilia show lack of NME5 protein expression and unchanged DNAH5 expression (Fig. 2 I, J).

**Figure 2:**
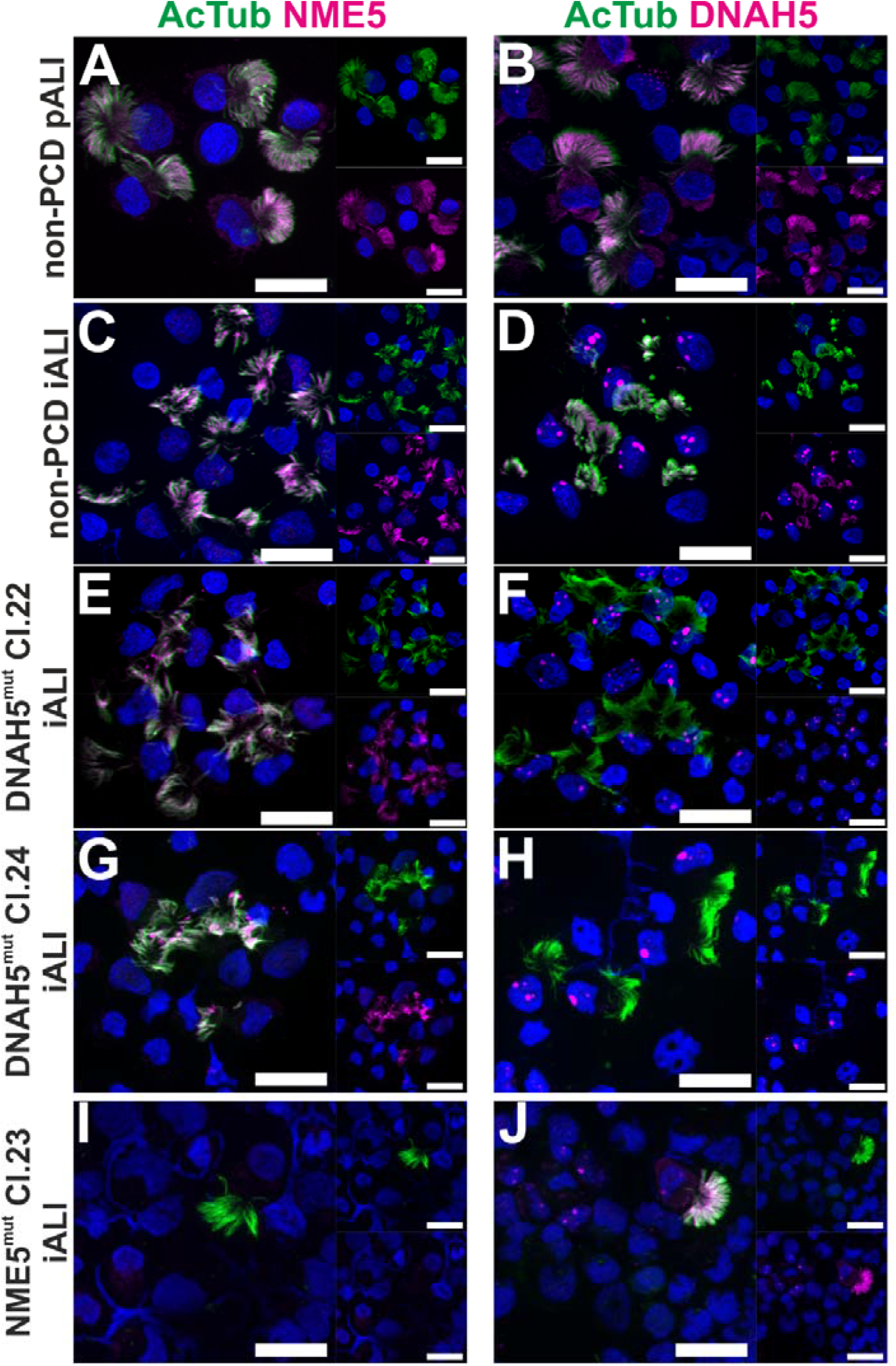
PCD specific hiPSC-derived epithelial cells show lack of ciliary protein expression of DNAH5 and NME5. Immunofluorescence staining of DNAH5 and NME5 of dissociated primary cell-derived (pALI) and hiPSC-derived ALI cultures (iALI) (day 26-31 after air lift) from different cell lines as indicated. Nuclei stained with DAPI (blue); scale bars represent 20 μm (A-J).

### Transmission electron microscopy analysis shows impact of DNAH5 and NME5 mutations on ciliary ultrastructure

To further investigate the impact of each specific mutation on the cilia axoneme ultra-structure, transmission electron microscopy (TEM) of generated iALI cultures was performed. Analysis of cilia of non-PCD diseased hiPSC-derived cells showed a normal 9+2 ultrastructure pattern in more than 95% of analyzed cilia axonemes (Fig. 3 A, B) as expected. In contrast, NME5-mutants show a 9+2 structure only in 56.6% of cilia axonemes and frequent numerical abnormalities or disorganization of cilia ultrastructure (Fig. 3 A, B). 5.1% of axonemes exhibited a 9+0 ultrastructure pattern, which is consistent with previous findings of Cho and colleagues [27]. Furthermore, we detected additional abnormal ultrastructural patterns. Most abundantly, 23.4% of axonemes exhibited an 8+1 pattern. A smaller subset of axonemes showed additional configurations of microtubule transpositions (e.g., 7+4, 8+4), additional microtubules (9+4 pattern) and/or microtubular disorganization. These findings resemble phenotypes associated with mutations in other radial spoke proteins (RSPH1, RSPH3) [28, 29].

**Figure 3:**
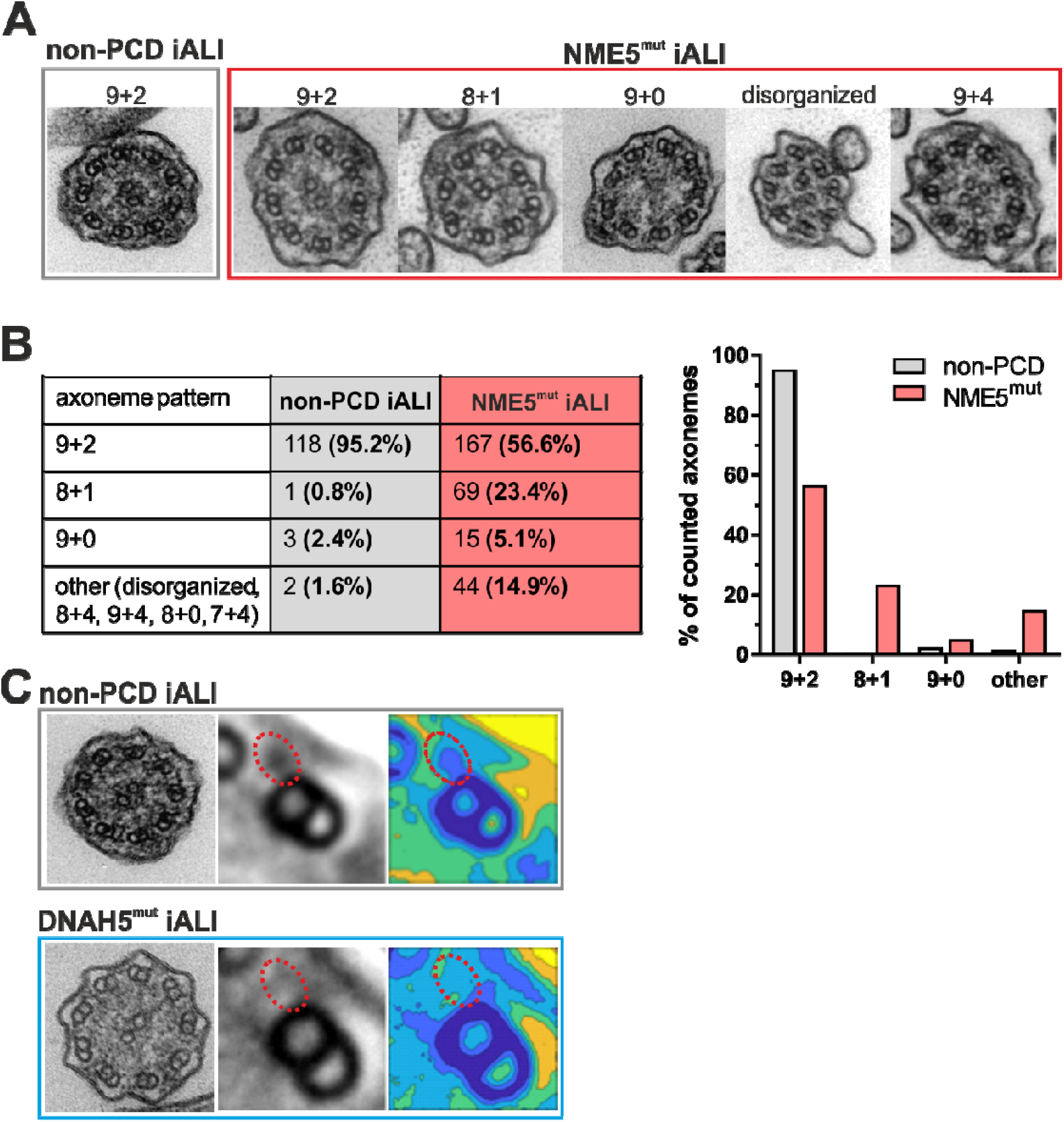
Ultrastructural defects in PCD hiPSC-derived ciliated epithelial cells can be detected by TEM. Representative transmission electron microscopy (TEM) images of cilia cross-sections from non-PCD and NME5^mut^ iALI-derived ciliated cells (A). Quantitative analysis of distribution of the most apparent axoneme patterns in non-PCD iALI and NME5^mut^ iALI axonemes (B). TEM image of non-PCD and DNAH5^mut^ Cl. 24 iALI cilia axoneme. Averaging of outer microtubule doublets (middle image) and generation of colour contour map of electron density (right image) was performed with PCD Detect software. Red dotted line presence (non-PCD) or absence (DNAH5^mut^) of outer dynein arms (C).

For investigating outer dynein arm (ODA) defects in DNAH5-mutated cilia, TEM cross-sections were analyzed using the PCD Detect software [30]. This software allows overlapping and averaging of microtubule features to generate enhanced signals and improved structural resolution. By averaging the electron density of multiple outer microtubule doublets, shortening of outer dynein arms in DNAH5-mutated cilia compared to non-PCD controls was detected and visualized in color coded density plots (Fig. 3 C).

### PCD specific hiPSC-derived ALI cultures show altered ciliary beat frequencies

To evaluate maturation and functional consequences of the DNAH5 and NME5 mutations in PCD, we measured ciliary beat frequencies (CBF) using high speed microscopy at day 35 after air lift or later. In non-PCD cells (pALI and iALI) ciliary beat was clearly visible directly in ALI cultures, enabling us to use tissue-level recordings and automated analysis of CBF (Fig. 4 A). In PCD cells (DNAH5^mut^ and NME5^mut^), ciliary beat was too sparse or abnormal for direct analysis of ALI cultures, and we therefore conducted manual analysis of CBF in single cell suspensions (Fig. 4 B). Non-PCD iALI cultures showed relatively low variation of CBFs and a median value of 8 Hz (IQR=1.3), which compared well to the similarly distributed CBF values in non-PCD pALI with a median value of 11.4 Hz (IQR=3.0). Compared to non-PCD iALIs, DNAH5^mut^ cultures showed significantly lower values of CBF, with a median of 2 Hz (IQR=1.2), whereas NME5 cultures exhibited significantly higher and highly variable values of CBF, ranging from 5 to 19 Hz with a median of 15 Hz (IQR=4.5) (Fig. 4 C).

**Figure 4:**
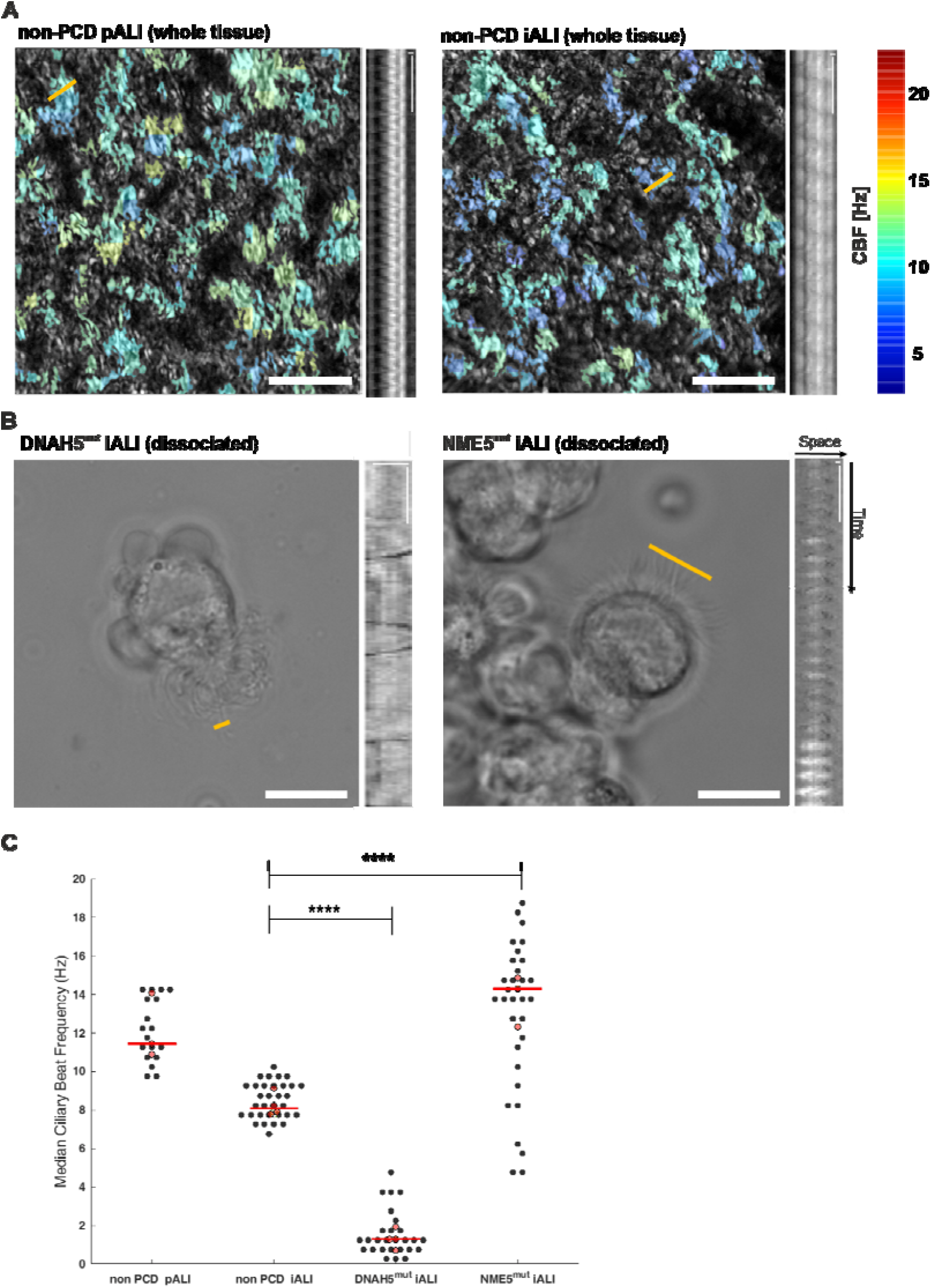
Ciliary beating frequencies are altered in PCD hiPSC-derived epithelial cells. Representative top-view heatmaps of ciliary beat frequencies in non-PCD pALI and non-PCD iALI cultures, color-coded by ciliary beat frequency and overlaid onto maximal projection of all movie frames. Scale bar: 50 μm. Associated kymographs for comparison. Scale bars kymographs: 1μm in space, 0.25 s in time (A). Representative image of DNAH5^mut^ and NME5^mut^ suspension movies and corresponding kymographs. Yellow line indicates position at which a kymograph was taken for CBF counting. Scale bar phase contrast images: 10um; Scale bars kymographs: 1μm in space, 0.25 s in time (B). Median of ciliary beat frequencies in all 4 cultures. Black dots: median value per box (non-PCD pALI and iALI), median value per FOV (DNAH5^mut^, NME5^mut^); red dots: median value across FOVs for each experiment (different isolation, differentiation or clone); red bar: median of all experiments. ^****^p<0.0001 (C)

### PCD specific hiPSC-derived ALI cultures show impaired mucociliary clearance (MCC)

Since functional MCC depends on many factors, including ciliary beat coordination and orientation [31, 32], CBF alone is not a good predictor of MCC [33]. Indeed, visually comparing the ciliary beat in suspended cells of all conditions revealed metachronal and coordinated beat in non-PCD pALI and iALI cultures (Suppl. Videos 1 and 2), whereas we observed sparse strokes and occasional vibrations in DNAH5-mutant cells (Suppl. Video 3), which is consistent with literature [23, 34, 35]. NME5-mutant cells showed an asymmetrical beating pattern (Suppl. Video 4) and sporadically exhibited circular instead of whipping cilia movements (data not shown). This has not been described for NME5 defects, yet, but resembles findings in other radial spoke defects [24, 36-38]. Altogether these defects suggest that the cilia beat in the PCD mutants might be ineffective at transporting fluids. We therefore proceeded to measure a direct proxy of MCC by recording the transport of suspended fluorescent microspheres on the intact tissue surface at day 35 post air lift or later (Fig. 5 A). As expected, the non-PCD pALI cultures generated the fastest flow that covered most of the tissue surface. Non-PCD iALI cultures created significantly faster and more expansive MCC than DNAH5 and NME5-mutated cultures (Fig. 5 B), demonstrating that the functionally healthy versus the diseased phenotype of mutated airway epithelia from PCD patients is exhibited by the hiPSC-derived cultures.

**Figure 5:**
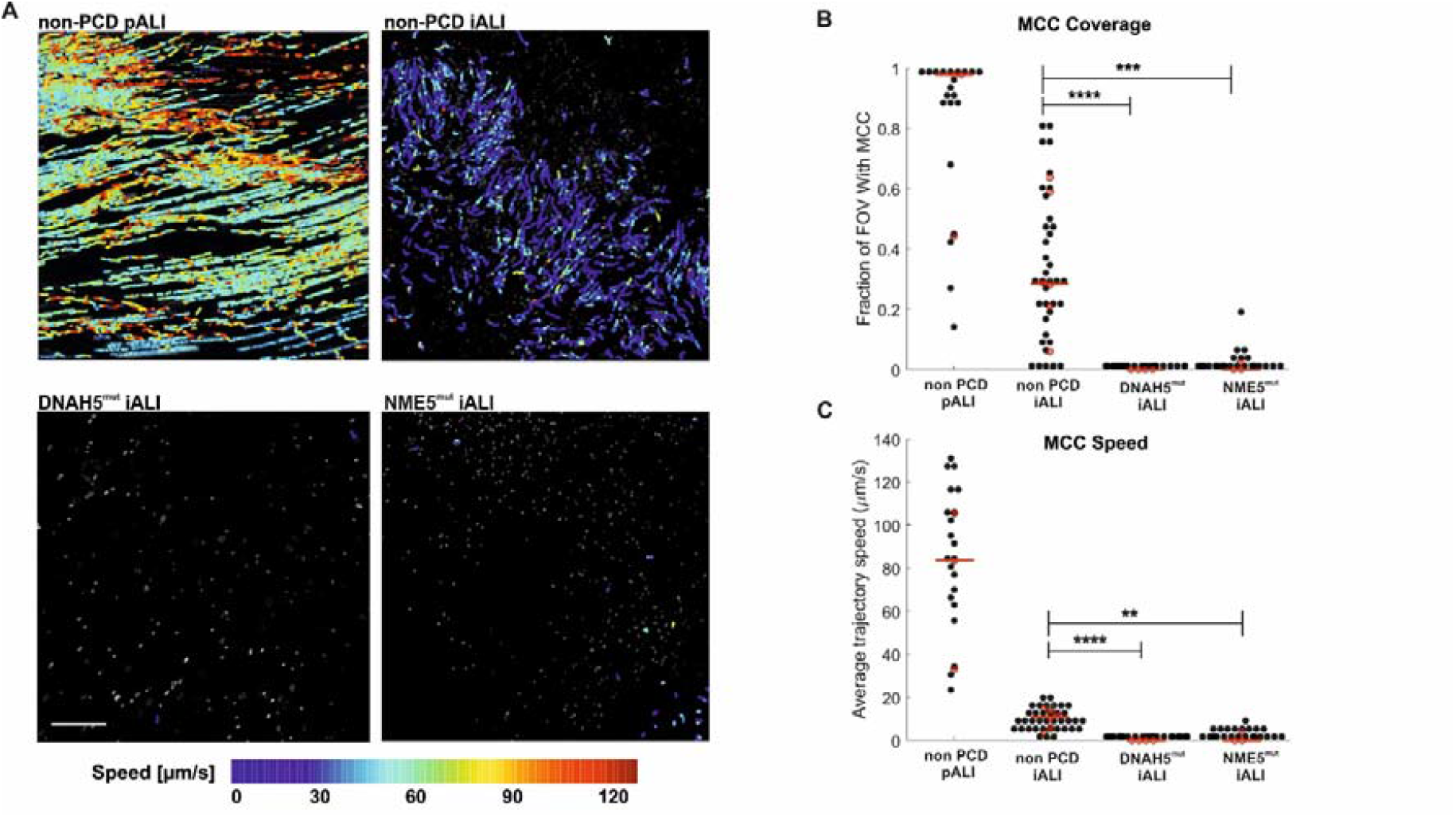
PCD specific hiPSC-derived ALI cultures show impaired mucociliary clearance. Representative plots of bead trajectories color-coded by instantaneous flow speed and overlaid onto maximal projection of all movie frames. Scale bar: 200 μm (A). Relative fraction of the field of view (FOV) with detected MCC (B). Average speed of flow trajectories (C). Black dots: average value per FOV; red dots: median value across FOVs for each experiment (different isolation, differentiation or clone); red bar: median of all experiments. ^**^p<0.01; ^***^p<0.001; ^****^p<0.0001 (C).

## 3. Material and Methods

### hiPSC Cultivation

All hiPSC lines ((non-PCD diseased (MHHi001-A) [25], DNAH5^mut^ (MHHi017-A (Clone 22) and MHHi017-B (Clone 24)) [10] and NME5^mut^ (MHHi019-B) [10]) were generated from CD34^pos^ cells using the CytoTune®-iPS 2.0 Sendai Reprogramming Kit (Thermo Fisher Scientific), following the manufacturer’s instructions. hiPSC were cultivated on Geltrex^™^ (Gibco) in E8 cell culture medium (in house-made). Medium was exchanged daily. Cells were passaged every 3 to 4 days using Accutase^™^ (Gibco) for up to 10 passages.

### Differentiation of hiPSCs Towards Lung Progenitor Cells

Seven days prior to the start of differentiation, hiPSCs were seeded at a low cell density (non-PCD: 1.2×10^4^ cells/cm^2^; NME5^mut^ and DNAH5^mut^: 2.0×10^4^ cells/cm^2^) in a Geltrex^™^ coated flask using E8 medium. Medium was supplemented with 10 μM Y-27632 (Tocris) for 24 hours (h) after seeding. Medium was exchanged daily. At day -4, preculture was initiated by supplementing E8 with Supplement 1, provided in the STEMdiff^™^ Definitive Endoderm Kit (STEMCELL Tech.). At day -1, cells were passaged using Accutase^™^ and seeded onto a new culture vessel (non-PCD: 0.33×10^4^ cells/cm^2^; DNAH5^mut^ Clone 22 (Cl. 22): 0.44×10^4^ cells/cm^2^; DNAH5^mut^ Cl. 24: 0.44×10^4^ cells/cm^2^, NME5^mut^: 0.44×10^4^ cells/cm^2^) in E8 supplemented with 10 μM Y-27632 and Supplement 1. After 24h, definitive endoderm differentiation was initiated by exchanging the medium to STEMDiff^™^ Basal Medium with Supplement CJ and MR. After 24h and 48h, medium was exchanged to STEMDiff^™^ Basal Medium with Supplement CJ. At day 3, cells were detached using Accutase^™^, analyzed for DE markers using flow cytometric analysis and plated for subsequent lung progenitor specification.

For anterior foregut specification, endodermal cells were plated at 12.5×10^4^ cells/cm^2^ for non-PCD; 21.9×10^4^ cells/cm^2^for DNAH5^mut^Cl. 22; 18.8×10^4^ cells/cm^2^for DNAH5^mut^ Cl. 24 and 31.3×10^4^ cells/cm^2^ for NME5^mut^ in basal media (Knockout DMEM (Gibco), 5% Knockout serum replacement (Gibco), 1% penicillin/streptomycin (Gibco), L-glutamine (Gibco), non-essential amino acids (Gibco) and 0.46 mM monothioglycerol (Sigma Aldrich)) supplemented with 3 μM Dorsomorphin (Sigma Aldrich), 10 μM SB431542 (provided by the Institute of Organic Chemistry, Leibniz University, Hannover, Germany) and 10 μM Y-27632 (anterior foregut media 1; AFE1). After 24h or 48h, media was exchanged to basal media supplemented with 2 μM IWP2 (Tocris), 10 μM SB431542 and 3 μM Dorsomorphin (only for WT and NME5^mut^) for 24 or 48h (anterior foregut media 2; AFE2). Conditions for optimal anterior foregut specification were determined in advance for each cell line individually and are listed in supplemental table 1. For induction of lung progenitor fate, medium was switched to basal medium with 10 ng/mL BMP4 (R&D Systems), 10 ng/mL FGF10 (R&D Systems) and 3 μM Chir99021 (provided by the Institute of Organic Chemistry, Leibniz University, Hannover, Germany) until day 14 of differentiation. Medium was exchanged daily. At day 14, lung progenitor cells were washed with phosphate-buffered saline without Mg^2+^/Ca^2+^ (PBS w/o) and dissociated using Accutase^™^ for 10-20 minutes (min) at 37°C. After purification using magnetic activated cell sorting, cells were plated onto transwells (Greiner Bio-One) for airway epithelial specification.

### Magnetic Activated Cell Sorting (MACS) of Lung Progenitors

At day 14 of differentiation, cells were detached from culture vessels as described above. Cells were harvested in basal media with 10 μM Y-27632 and spun at 216× g for 3 min at 4°C. After straining and counting, up to 15×10^6^ cells were first incubated in MACS buffer (PBS w/o (Gibco), 1.0% BSA (Sigma Aldrich), 2 mM EDTA (Sigma Aldrich)) with FcR blocking (Miltenyi Biotec) for 10 min at 4°C and mouse anti-human CPM antibody (FUJIFILM Wako; 1:200 dilution) for additional 20 min at 4°C. Unbound antibody was removed by washing once with MACS buffer, before cells were incubated in anti-mouse IgG 2a+b microbeads (Miltenyi Biotec) for 20 min at 4°C. After washing once with MACS buffer, cells were separated using the QuadroMACS Separator (Miltenyi Biotec). NKX2.1 flow cytometry analysis was used to determine the proportion of lung progenitor cells before and after sorting.

### Flow Cytometry Analysis

For assessment of successful DE induction, flow cytometry analysis was performed at day 3 of differentiation according to our previously published protocol [39, 40]. Shortly, cells were detached with Accutase^™^ and stained with anti-CXCR4, anti-cKIT and anti-EpCAM in FACS Buffer (PBS w/o, 1.0% FCS (Pan Biotech), 1 mM EDTA) for 30 min on ice. After washing twice with FACS buffer, cells were resuspended in FACS buffer containing 1.7 μg/mL DAPI (Sigma-Aldrich) for dead cell exclusion for flow cytometry measurement. Lung progenitor cultures at day 14 of differentiation were detached as described above and fixed and permeabilized using the transcription factor staining buffer kit (Miltenyi Biotec). After washing with flow buffer (PBS w/o containing 1.0% BSA), cells were stained with anti-NKX2.1 for 30 min on ice. Cells were washed three times with FACS buffer and resuspended in FACS buffer for flow cytometry measurement.

Antibodies and respective dilutions are listed in supplemental table 3. All flow cytometry measurements were performed using the MACSQuant Analyzer 10 (Miltenyi Biotec) and data were analyzed with FlowJo software (Ashland, OR).

### Differentiation of Lung Progenitor Cells Towards Pseudostratified Airway Epithelium on Air-Liquid Interface Culture

After MACS sorting, purified lung progenitor cells were resuspended in small airway epithelial cell growth medium (SAECGM; PromoCell) supplemented with 1% penicillin/streptomycin (Gibco), 1 μM A83-01 (Tocris), 0.2 μM DMH-1 (Tocris) and 5 μM Y-27632 [41]. For initiation of air-liquid interface (ALI) culture, 2.65×10^5^ cells/cm^2^ were seeded per transwell (Greiner Bio-One), coated with 804G conditioned medium [41]. During expansion phase, medium was renewed every other day in apical and basal chamber. Once cells have formed a confluent cell layer (usually after 4 days), medium in apical and basal chamber was switched to PneumaCult^™^-ALI medium (STEMCELL Tech.) containing 1% penicillin/streptomycin. After 48h, medium in the apical chamber was removed to conduct the air lift and medium in the basal chamber was renewed. Cells were differentiated on ALI cultures for 28 days for molecular characterization and >35 days for functionality assessment with medium exchanges every two to three days.

### Quantitative RT-PCR

ALI cultures were lysed at day 28 after air lift using TRIzol (Thermo Fisher Scientific). RNA isolation was performed using the Nucleospin RNA II Kit (Macherey-Nagel) and reverse transcription of 500 ng RNA was done using the RevertAid H Minus First Strand cDNA Synthesis Kit (Thermo Fisher Scientific). cDNA was diluted 1:5 and RT-qPCR was done using SsoAdvanced^™^ Universal SYBR Green Supermix (Bio-Rad) with PrimePCR Assays (CCDC40, CK5) or self-designed primers (FOXJ1, P63, NGFR, NKX2.1, MUC5AC, CCSP, betaActin, GAPDH). Primer sequences and assay IDs are listed in supplementary table 2. Samples were pipetted as duplicates and run on the CFX Connect^™^ Real-Time System (Bio-Rad). Ct values were averaged, normalized to the housekeeping genes GAPDH and betaActin.

### Cytospin

Prior to staining for ciliary protein markers, airway epithelium was dissociated to single cells or small clumps. For this purpose, the membrane of day 28 ALI cultures was removed from the transwell using a scalpel and incubated in Accutase^™^ supplemented with 200 μg/mL DNase I (Roche) for 15-20 min in a shaking 37°C water bath. Dissociation was stopped with RPMI medium (Gibco) and cells were spun at 138x g for 3 min. After resuspending in RPMI medium, cells were centrifuged onto a Superfrost® microscope slide at 650 rpm for 5 min using Cytospin2 (Shandon). Slides were dried at room temperature and stored at −20°C until used.

### Paraffin Embedding and Sectioning

For fixation, medium was aspirated from apical and basal side and replaced with 4% parafomaldehyde (PFA) and was incubated for 15 min at room temperature. In order to remove mucus from the airway epithelium, the apical side of ALI cultures was washed repeatedly with PBS (Gibco) before fixation. After fixation, transwells were washed with and stored in PBS at 4°C. For embedding, membrane was cut out of the transwell using a scalpel, dehydrated (Leica Biosystems) and embedded in paraffin (Leica Biosystems). Using a microtome (Leica Biosystems), cross-sections of 3 μm thickness were generated.

### Immunofluorescence staining

Before staining paraffin embedded samples, paraffin was removed from cross-sections by first heating the sections to 60°C for 30 min, followed by incubation in xylol for 2× 15 min. Rehydration of the sections was done by stepwise incubation in ethanol (100%, 90%, 80% and 70%) for 2 min each and finally, incubation in water. For antigen retrieval, slides were incubated in 10 mM citrate buffer, pH6 (Sigma Aldrich) for 60 min at 133 °C.

Prior to staining of cytospin samples, slides were taken from the −20°C storage and fixed immediately with 4% PFA for 2 min at room temperature.

For immunofluorescence staining of cytospin samples and cross-sections, samples were washed twice with Tris-buffered saline (TBS) and were blocked with blocking buffer (TBS containing 5% donkey serum (Biozol) and 0.25% Triton X-100 (Sigma-Aldrich)) for 1h at room temperature. Primary antibody incubation was performed in staining buffer (TBS with 1% BSA) at 4°C overnight. Primary antibodies and respective dilutions are listed in supplemental table 3. Slides were washed three times with TBS and secondary antibody incubation was performed in staining buffer for 30 min at room temperature. Secondary antibodies are listed in supplemental table 4. Samples were washed three times with TBS and nuclei were stained with TBS containing 1.7 μg/mL DAPI (Sigma-Aldrich) for 5 min. After washing three more times with TBS, sections were mounted with mounting medium (Dako) and dried over night at room temperature. Images were taken with the Axio Observer A7 (Zeiss) by optical sectioning using the Apotome. Pictures were processed using ZenBlue 3.0 software (Zeiss).

### Transepithelial Electrical Resistance (TEER) Analysis

Prior to measurement, ALI cultures were supplied with fresh PneumaCult^™^-ALI medium on the basal side and covered with 750 μl of PBS supplemented with 100 μg/mL Primocin (InvivoGen) on the apical side of the transwell. Electrical resistance between apical and basal chamber is measured with an EVOM3 electrical volt-ohm meter (World Precision Instruments) using the STX4 electrode (R_Total_). Measurements were done in technical duplicated. A transwell without cells was used for blanking (R_Blank_). TEER values were calculated as follows:

(1) R_Tissue_ [Ω] = R_Total_ – R_Blank_

(2) TEER [Ω × cm^2^] = R_Tissue_ [Ω] × surface area [cm^2^]

### Transmission Electron Microscopy

ALI cultures were washed with PBS as described above and subsequently fixed and stored in 150 mM HEPES buffer containing 1.5% PFA and 1.5% glutaraldehyde. Membranes were cut off the ALI inserts and fixation and embedding was done as described in [42]

### PCD Detect Software Analysis

For assessment of ultrastructural dynein arm defects, PCD Detect Software was used, following the developers’ workflow and using the same alignment settings [30]. From each cilia cross-section, 9 cut-outs, each including a microtubule doublet with outer and inner dynein arms, were manually selected. At least 5 cilia cross-sections (equivalent to 45 cut-outs) from different cells were analyzed in each sample. In this study we present the averaging of all features as color contour maps of electron density (blue: high density; yellow: low density).

### Mucociliary Clearance (MCC) Measurement

Only cultures at day 35 of ALI or older were used for MCC measurements to ensure substantial maturity of MCC. To remove accumulated mucus and debris, ALI cultures were submerged with PBS for 10 minutes, then the PBS was carefully suctioned off. To visualize MCC, fluorescent microspheres (1 micron diameter, Invitrogen) were diluted 1:1000 in PBS, and 20 μl of this suspension were added to the surface of each ALI culture. Particle movement was recorded using a Zeiss Axioscope fluorescence microscope equipped with an Orca Flash 4.0 camera (Hamamatsu) and a temperature-controlled chamber that was preheated to 37° C. Apart from a few initial test recordings, standardized acquisition settings using a 10× objective were 1024×1024 pixels with 1.3 μm/pixel resolution at a framerate of 20 frames per second for a total duration of 10 seconds. For each condition, at least 3 experiments (different differentiations or clones (hiPSCs), or different isolations (pALI)) were performed. In each experiment, 5 fields of view (FOVs) with visible particle movement (sometimes only Brownian motion) were recorded from 2 insert cultures each. The particle trajectories were extracted from the raw movies using ImageJ/Fiji [43] with the Trackmate plugin [44]. Movies that did not contain analyzable bead signals due to debris or overgrowth were excluded. Trajectories were further analyzed to compute the average trajectory speed Uk and MCC coverage C_k_ (the fraction of the FOV containing MCC) of each FOV k using customized code in Matlab (Mathworks), as reported previously [33]. For statistical analysis, the pooled FOV-averaged trajectory speed and MCC coverage values (the U_k_ and C_k_ for all k = 1…n FOVs, respectively) per condition were compared with the non-parametric Kruskal– Wallis test, followed by Tukey-Kramer multi-comparison test, to test for different median values.

### Ciliary Beat Frequency (CBF) Measurement

Only cultures at day 35 of ALI or older were used for CBF measurements. CBF was recorded using a Zeiss Axioscope microscope, allowing for oblique phase contrast recordings, equipped with an Orca Flash 4.0 camera (Hamamatsu) and a temperature-controlled chamber that was preheated to 37° C. Koehler illumination was set up for all recordings. Two different procedures were performed to measure CBF. In non-PCD pALI and non-PCD iALI cultures CBF was measured at the tissue level, and single-cell CBF was measured in DNAH5^mut^ and NME5^mut^ cultures, as the movement of cilia could not be resolved at the tissue level. For tissue-level CBF measurements, ALI cultures were washed as above. High speed movies of CBF (ca. 140 frames per second for a duration of at least 1.5 seconds) were recorded using a 40× long distance phase contrast objective at a spatial resolution of 0.3 μm/pixel in a 512×512 pixel frame. For each condition, at least 3 experiments (different differentiations (hiPSCs) or different isolations (pALI)) were performed. From cultures of each experiment, 6 FOVs with visible CBF were recorded. We measured cilia beat frequencies by applying Fourier spectral analysis, as previously described [45]. The mean CBF was calculated for each window of 32×32 pixels, resulting in a maximum of 16 mean CBF values per FOV if all windows contained ciliary beat, and from these values the median CBF of the entire FOV was determined.

For single-cell CBF measurements, the cell layer was dissociated by incubating the ALI cultures for 20 min in prewarmed Accutase^™^ in a conical tube. DNase I at a concentration of 200 μg/mL was added to remove any free DNA resulting from the dissociation process. After 20 min warm culture medium was added at a ratio of 3:1 and the cells were centrifuged at 170 to 240 ×g for 5 min. The resulting pellet was resuspended in 1 mL culture medium to obtain a high cell density and 10 μl of this suspension was placed onto a glass bottom imaging dish (Ibidi). To confirm viability of the cells, the same amount of life-dead stain (Invitrogen) was added to the cell suspension droplet and the solution was covered with a round glass coverslip in order to prevent evaporation. The measurements were performed using a 40x oil phase contrast objective (NA 1.4) under standardized acquisition settings, which were 800 frames per second for a duration of 2 seconds (NME5^mut^) and up to 10 seconds (DNAH5^mut^) of recording time and a spatial resolution of 0.16 μm/pixel in a 265×265 pixel frame. For each condition, at least 3 experiments (different differentiation (hiPSCs) or different isolations (pALI)) were performed, and in each experiment at least 9 FOVs, i.e., 9 ciliated cells, were recorded. After each CBF movie recording a brightfield image and fluorescent images of the life-dead staining were recorded to confirm viability. The recordings were analyzed using the kymograph function in ImageJ. Kymographs of individual or groups of cilia were generated and the frequency was calculated by counting the amount of pixel intensity changes over time in the kymograph. Whenever a wave pattern was observed, the number of peaks were counted, else the dark lines were counted and divided by two to obtain a full beat cycle count. In the case of non-harmonic oscillation observed in DNAH5^mut^-derived ciliated cells, stationary vibrations were not included into the count. A median was calculated from all counted frequencies per ciliated cell. For statistical analysis, the pooled tissue-level medians and single-cell medians per condition were compared with the non-parametric Kruskal–Wallis test, followed by Tukey-Kramer multi-comparison test, to test for different median values.

## 4. Discussion

To date, mainly primary cells or non-airway immortalized cell lines (e.g., HEK293 cell line) have been used for PCD *in vitro* modelling or drug screening [46-48]. However, both cell sources harbor severe limitations for such applications. Non-airway immortalized cell lines do not allow assessment of therapeutic effect on a functional level, and primary airway cells, while they represent the PCD phenotype well, are limited in their accessibility and expandability. Even though it has been demonstrated that primary cells can be expanded over 35 passages and maintained their differentiation potential [49], this requires laborious matrix embedded cultures, which limits the cell yield. Furthermore, targeted gene editing is not possible on a clonal level. hiPSCs exhibit infinite self-renewal and can be differentiated into the cell type of interest, thereby overcoming limitations of primary cells. Moreover, PCD-patient specific hiPSCs can be used to study the disease on different disease implicated tissues (airway cilia, sperm cells, nodal cilia) and can be genetically modified on a clonal level, e.g., for introducing reporter systems for screening readout. Hence, hiPSCs represent a valuable tool for PCD disease modeling and screening applications.

Current differentiation protocols for generation of airway epithelium from hiPSCs often include an intermediate 3D culture step, in which lung progenitor cells are embedded in Matrigel droplets, prior to differentiation to airway epithelium [25, 26]. Hence, these protocols are lengthy (28 -46 days until initiation of ciliation by air lift or supplementation with DAPT [25, 26]) and scale up of cell production is severely limited due to the small scale and laborious 3D culture. Here, were describe an efficient airway differentiation protocol that takes only 20 days until initiation of ciliation. Moreover, our protocol is not dependent on complex Matrigel-dependent organoid culture steps, hence more defined and scalable for future high throughput drug screening applications.

Sone et al. previously reported a hiPSC-based disease model of PCD-causing mutations in *DNAH11, HEATR2* and *PIH1D3* using a complex airway-on-a-chip technology [24]. By applying fluid shear stress via defined cell culture medium flow rates on the apical side of the forming epithelium, planar cell polarity can be effectively induced in ciliated cells. Hence, compared to simple transwell ALI cultures, the chip technology allows the application and investigation of mechanical effects on the cells. However, such culture-on-a-chip is technically complex, expensive and also harbors several limitations. The microfluidic design of the system hampers the establishment of chip cultures in unexperienced labs, the access to the cultured epithelium, the application of standard characterization methods (e.g. TEM, staining of cross-sections) and functional high throughput readouts using plate reader technology. In particular, electrophysiological assessments such as TEER and Ussing chamber measurements, which are valuable techniques to determine epithelial quality and functionality [27, 50], are not feasible in commercially available airway chips. Integration of electrodes is possible in custom-made chip systems but this approach is very laborious and requires microfabrication infrastructure and expertise [51, 52]. Moreover, the submerged culture system utilized by Sone and colleagues [24] does not allow to investigate the implication of mucus properties in PCD disease development and infection processes. Additionally, for future compound screening applications, downscaling of culture size to high or medium throughput formats is indispensable and not yet possible for chip cultures.

In contrast, transwell ALI cultures are compatible with standard readouts, easier to establish due to the commercial availability and handling simplicity, and available in various formats including 96-well plates (Corning #100-0419), allowing higher throughput screening of pharmacological compounds. Altogether, the airway-on-a-chip system is a valuable tool to investigate specific research questions under dynamic flow conditions; however, the transwell ALI culture system remains indispensable to be used as a complementary system, in particular for screening purposes.

In the present study, we describe the establishment of a hiPSC-based PCD *in vitro* disease model using patient lines containing mutations in the most frequently mutated and well described PCD-associated gene *DNAH5* [34, 35], and in the recently as PCD-associated described gene *NME5* [27], respectively. Our data show that mutated *DNAH5* results in loss of DNAH5 protein, altered ODA ultrastructure and reduced CBF, consistent with previous published iPSC-based models [23]. Additionally, we investigated bead transport on the tissue level that displayed significantly reduced MCC capacity in *DNAH5*-mutated cultures. Moreover, we demonstrate the application of the newly developed PCD Detect software [30] that allows to identify ODA defects. Previous disease models mainly focused on describing a severe DNAH5 ultrastructural phenotype that exhibits total absence of ODA [23], but mutated *DNAH5* can also result in less prominent ODA defects such as dynein arm truncation [4]. Reliable analysis of the whole range of ODA phenotypes is necessary for diagnostics and experimental readouts; however, analyzing ultrastructure with only subtle defects can be challenging and strongly rely on the investigators expertise [30]. Hence, we used the PCD Detect software as an investigator-independent and time efficient diagnostic method [30] to identify subtle ODA shortening in our hiPSC *DNAH5*-mutated cultures. Altogether, the detailed characterization of our *DNAH5*-mutated iALI cultures serves as a prove-of-concept to demonstrate that our *in vitro* model recapitulates the *DNAH5*^*mut*^ -specific phenotype and can be captured in detail by utilizing improved readout methods.

Next, we aimed at modelling the impact of mutations in the gene *NME5*, which was recently identified as PCD-associated. NME5 is part of the radial spoke neck; however, so far, only little is known about the function of NME5 and the pathomechanism of its mutation [27, 53]. Solely Cho and colleagues studied the effect of mutant NME5 in humans and have described an abnormal axonemal ultrastructure in mutant ciliated cells (loss of the central pair; 9+0 pattern) [27]. Here, we investigated the effects of mutated *NME5* in more detail. Our TEM results are consistent with the previously reported loss of central pair (9+0) in *NME5*-mutated cilia, and moreover show additional ultrastructural abnormalities, such as outer doublet transposition (8+1, 7+4, 8+4) and presence of additional microtubules (9+4), which have not been described for this mutation before. Importantly, similar ultrastructural abnormalities were reported in other radial spoke protein defects [22, 38, 54-56]. These findings suggest that NME5 is indispensable for proper radial spoke assembly and stabilization of the central pair complex.

To further investigate the effect of mutated *NME5* on single cell and tissue level, we performed high speed video microscopy (HSVM). Single cell functional analysis revealed that *NME5-*mutant ciliated cells beat in an uncoordinated way and exhibited partially circular cilia movements that resembled 9+0 nodal cilia beating. These nodal-like beat kinematics have been previously reported to be a consequence of central pair and transposition defects [24, 36-38]. Additionally, *NME5*-mutated ciliated cells showed an increased variance of CBFs, compared to controls. By assessing the mucociliary transport capacity, we could additionally demonstrate that these alterations lead to a significantly diminished mucociliary clearance capacity (coverage and speed) compared to non-PCD controls (iALI and pALI). Altogether, these findings indicate that mutated *NME5* not only impairs beating synchrony and pattern of a given cell, but it also potentially prevents tissue-level metachronal wave formation due to divergent beat frequencies, which breaks down mucus clearance [57].

Notably, also non-PCD iALIs showed reduced MCC compared to primary cell controls (non-PCD pALI), despite comparable CBF parameters (median and variance) and synchronous cilia beating. This heterogeneity may have been neglected in other studies that used smaller fields of view for measuring bead transport; e.g., Sone et al. used a 100 μm × 100 μm fields of view (FOV) [24] compared to our 600 μm × 600 FOV. Our larger FOV allows us to obtain a more representative picture of the tissue, which might be particularly necessary to detect phenotypes on a tissue level (as in the case of *NME5*^*mut*^ cells). Further underlying causes for the observed differences of pALI and iALI cultures may include an insufficient maturation of the iALIs. Despite analyzing the cultures earliest at day 35 after air lift, maturation of ciliary beating (synchrony, force generation and cilia length) might be not fully completed. So far, the maturation of cilia beating during differentiation is not well understood, and investigation of bead transport over the course of differentiation and analysis of cilia length remains to be done for further understanding. In addition, the higher heterogeneity of hiPSC-derived cultures (ciliated cells appearing in clusters and/or uneven epithelium surface) compared to pALIs might reduce MCC coverage and average trajectory speed, since bead transport slows down in non-ciliated areas.

Taking together, here we present a hiPSC-based PCD model platform utilizing a simple air-liquid interface culture system that recapitulates mutation specific phenotypes by detailed characterization on a molecular (mRNA, protein, ultrastructure) and functional (CBF, HSVM, bead transport) level. In addition, our system allows down-scaling of the culture format and up-scaling of cell yield, which are essential prerequisites for upcoming higher throughput screening applications.

## Author Contributions

“Conceptualization R.O., F.C.R. and U.M.; software, D.R., J.N. B.O.S.; formal analysis, D.R., J.N. L.v.S., J.H..; investigation, L.v.S., D.P., N.C., J.Z..; data curation, L.v.S., J.H.; writing—original draft preparation, L.v.S., J.N. D.R., F.C.R, R.O., U.M..; visualization, B.O.S., J.H., L.v.S. D.R., J.N..; supervision, U.M., R.O..; project administration, U.M., R.O.; funding acquisition, U.M., R.O, F.C.R. All authors have read and agreed to the published version of the manuscript.”

## Funding

“This research was funded by German Center for Lung research (DZL; 82DZL002A1)

## Notes

### Competing Interest Statement

The authors have declared no competing interest.

